# Large-scale cellular-resolution imaging of neural activity in freely behaving mice

**DOI:** 10.1101/2021.01.15.426462

**Authors:** D.P. Leman, I.A. Chen, K.A. Bolding, J. Tai, L.K. Wilmerding, H.J. Gritton, Y. Cohen, W.W. Yen, L.N. Perkins, W.A. Liberti, K. Kilic, X. Han, A. Cruz-Martín, T.J. Gardner, T.M. Otchy, I.G. Davison

**Affiliations:** Department of Biology, Boston University; Department of Biomedical Engineering, Boston University; Department of Psychological and Brain Sciences, Boston University; Neurophotonics Center, Boston University; Department of Electrical Engineering and Computer Science, University of California at Berkeley; Knight Campus, University of Oregon; Department of Biology, Brandeis University; Department of Comparative Biosciences, University of Illinois at Urbana-Champaign; Department of Neurosurgery, Massachusetts General Hospital

## Abstract

Miniaturized microscopes for head-mounted fluorescence imaging are powerful tools for visualizing neural activity during naturalistic behaviors, but the restricted field of view of first-generation ‘miniscopes’ limits the size of neural populations accessible for imaging. Here we describe a novel miniaturized mesoscope offering cellular-resolution imaging over areas spanning several millimeters in freely moving mice. This system enables comprehensive visualization of activity across entire brain regions or interactions across areas.

## Main

Freely moving animals engage in many complex, ethologically relevant behaviors that are difficult to faithfully reproduce in head-fixed animals, ranging from social interactions to navigation. Miniaturized head-mounted systems for fluorescence imaging have made it possible to visualize the neural activity patterns underlying such behaviors^1–4^, and evaluate how they are shaped by learning over days to weeks^5–8^. While revealing new insights into the links between brain and behavior, first-generation systems typically offer imaging areas on the order of 1 mm^2^ or smaller^1, 9^, constraining the size of the neural populations that can be monitored and largely precluding measurements spanning multiple brain regions, which currently requires multiple implants^10, 11^.

While mesoscale systems for imaging areas up to several millimeters across have become available^12–16^, current tabletop designs require physical restraint, prohibiting many naturalistic behaviors demanding freedom of movement^5, 7, 17^. We set out to engineer a miniaturized, head-mounted mesoscale system for comprehensive, cellular-resolution imaging across extended brain areas. Our design goals included (i) field-of-view of >3 x 3 mm^2^, capable of encompassing multiple brain areas; (ii) optical resolution ≤5 μm for single-cell measurements; (iii) high numerical aperture (NA) for high efficiency and contrast; (iv) minimal size and weight for use in mice and other small animal models; and (v) economical off-the-shelf components to facilitate adoption by the community.

Building on our established conventional miniscope platform^9^, we expanded field-of-view by replacing the conventional gradient-index objective with a series of 6 mm diameter lenses providing ~1.3X magnification (Fig. 1a). Off-the-shelf optics were assembled in a 3-D printed housing (FormLabs Form2) that also holds the LED excitation source, collimation optics, and associated filters (Fig. 1b; see Supplementary Methods). We used an optical configuration that maximizes balance and freedom of movement, where excitation is delivered from above with a 470 nm LED, and two short-pass dichroic mirrors reflect emitted fluorescence signals towards the rear of the housing and then upwards onto the horizontally oriented CMOS sensor. Collection efficiency and resolving power are set by an aperture stop between the lens groups (Fig. 1a), which can be sized during printing to enhance efficiency at the expense of off-axis resolution and vice versa. The mesoscope has a working distance of ~2.5 mm, captures a field-of-view of approximately 4 mm X 4 mm (Fig. 1c), and resolves ≥150 lines/mm in the center of the imaging field when paired with a 2.2 μm pixel CMOS sensor (Fig. 1d; using 3 mm aperture/0.15 NA). Individual neurons were readily visualized in fluorescently labeled tissue sections (Fig. 1e,f), and the system was carried by mice with little difficulty (weight, 4.2-4.5 grams; housing size, 17 mm x 36 mm x 9 mm). Overall, our design opens up new possibilities for capturing cellular-scale activity across extended brain networks during complex behaviors.

**Figure 1.**
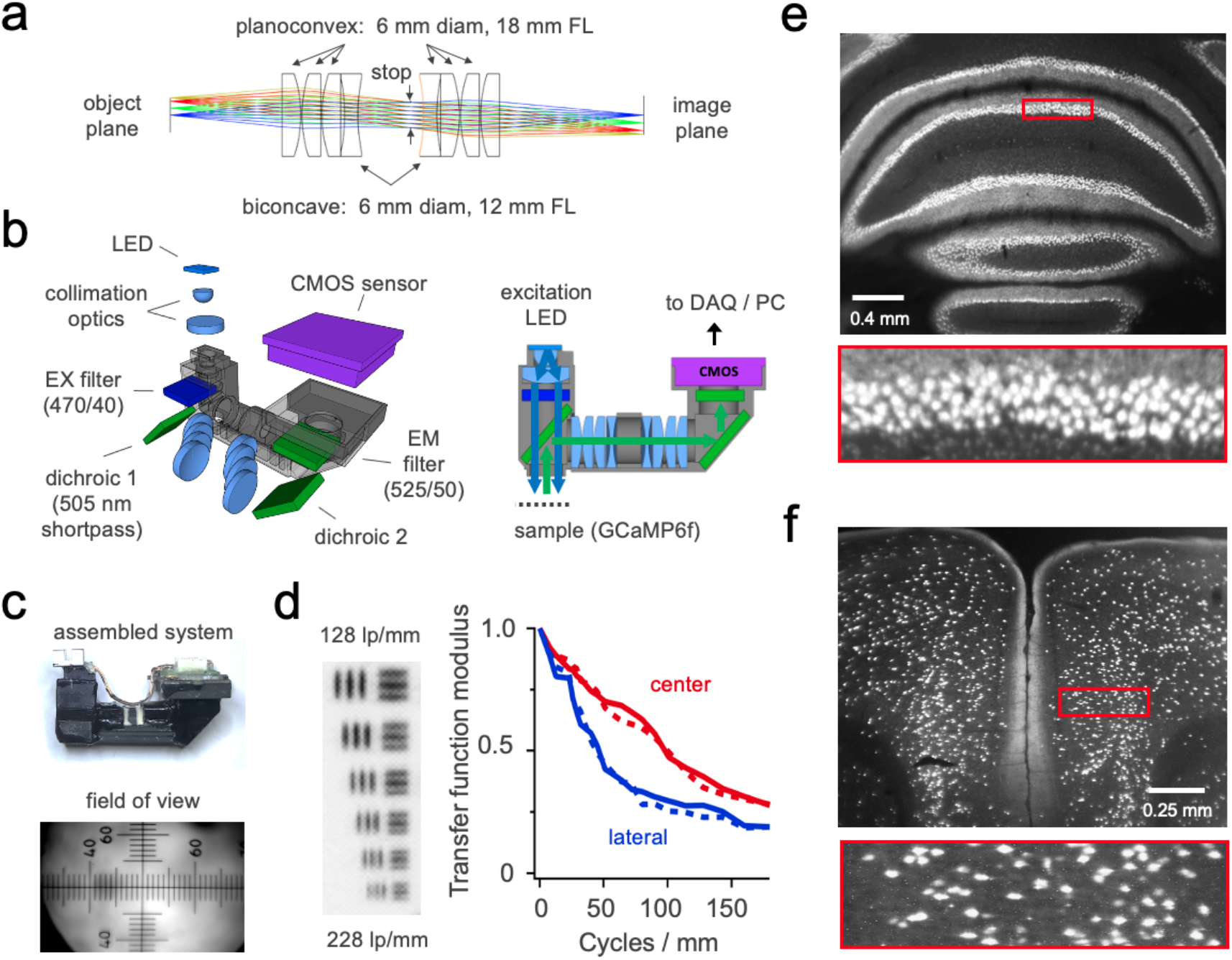
Mesoscope design and characterization. **(a)** Configuration of the optical pathway. **(b)** Left, exploded view showing system components. Right, schematic of the illumination and collection pathways. **(c)** Assembled mesoscope and image of ruled micrometer showing dimensions of the imaging area (~4 mm X 3 mm; small ticks, 100 μm; largest tick, 1 mm. **(d)** Characterization of optical performance. Left, segment of resolution target imaged with the mesoscope. Line density varies from 128 to 228 line pairs / mm (7.8 to 4.3 μm spacing, top and bottom respectively). Right, measured optical transfer function in both horizontal (solid) and vertical (dashed) dimensions. **(e,f)** Mesoscope images of ZsGreen-labeled interneurons in fixed samples of cerebellum (top) and cerebral cortex (bottom), illustrating single-cell resolution.

To apply the mesoscope for visualizing neural activity in freely moving mice, we began with widefield imaging of sensory representations in main olfactory bulb (MOB), where odor cues are encoded by unique activity patterns distributed across spatially segregated sensory pathways^18^. While extensive work has characterized how odor identity is encoded in immobilized animals^18, 19^, little is known about the sensory signals that guide active, spatially directed behaviors such as navigation and localization of odor cues^20^.

We used the expanded field of view to capture activity across both hemispheres of MOB as mice actively explored odor sources dispersed throughout a rectangular arena (Fig. 2a). Time-averaged sensory maps elicited by natural investigatory behaviors were consistent with previous work in head-fixed animals^18, 19^ (Fig. 2b). However, image series acquired at 30-60 Hz revealed rich dynamics that varied with proximity to the odor source as well as timevarying sniff patterns (Fig. 2c,d; Supplemental Video 1). Notably, the left and right MOB hemispheres often showed robust differences in the intensity and timing of activity, suggesting that even single-sniff sensory samples contain ‘stereo’ odor information that may inform navigational decisions^21^ (Fig. 2e; similar results obtained from 5 additional mice). Thus, mesoscopic imaging allows large-scale measurements of sensory dynamics during naturalistic, free-moving behaviors that are inaccessible in head-fixed experiments.

**Figure 2.**
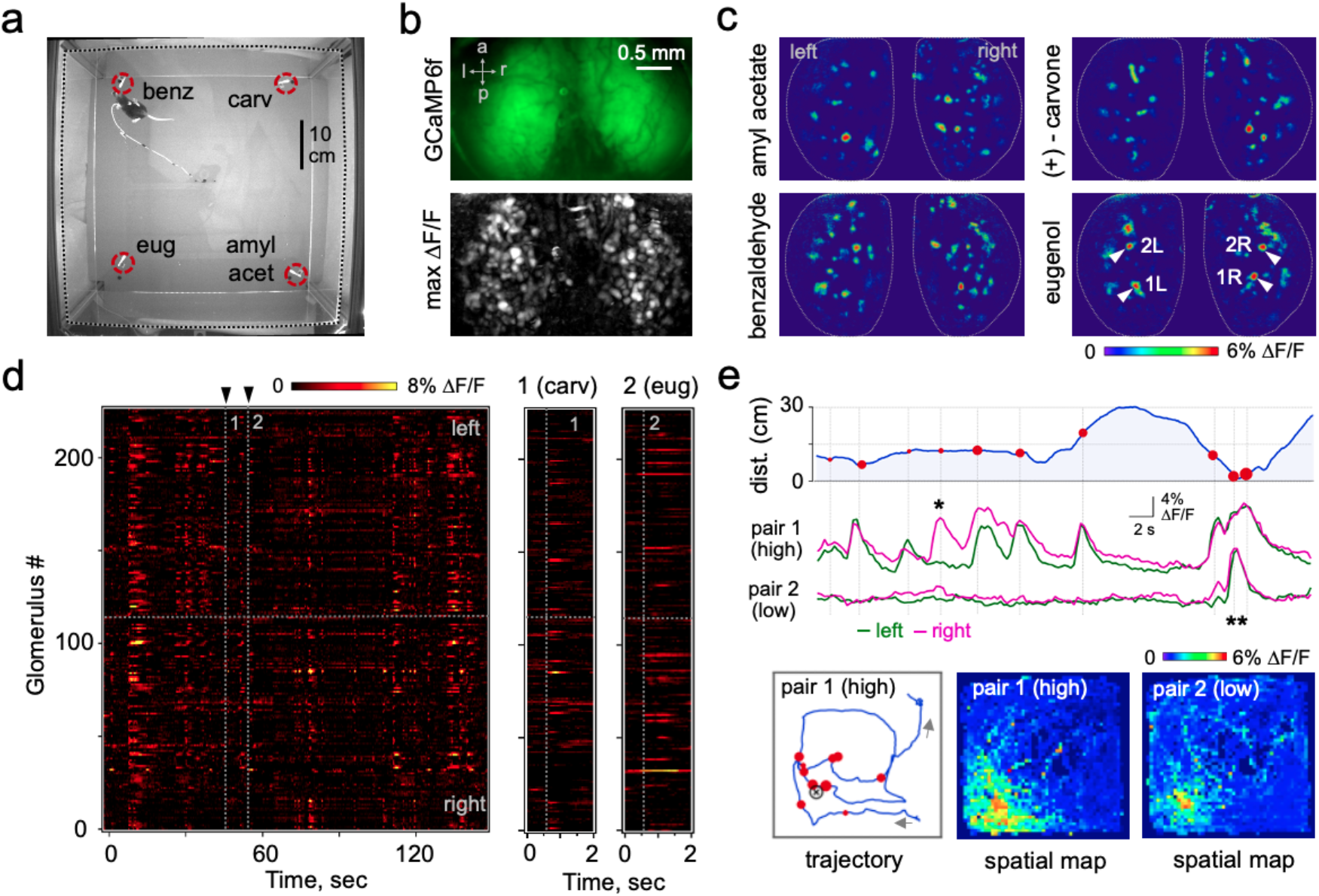
Bilateral imaging of olfactory dynamics during natural odor investigation. **(a)** Bottom view of behavioral arena. Dashed square indicates walls; circles, odor sources (benz, benzaldehyde; carv, (+)-carvone; eug, eugenol; amyl acet, amyl acetate). **(b)** Top, false-color image of raw GCaMP6f fluorescence in dorsal MOB. Bottom, maximum projection of the ΔF/F timeseries showing discrete ROIs corresponding to sensory glomeruli. **(c)** Time-averaged maps of sensory responses showing distinct activity patterns for 4 different odorants. Arrowheads indicate glomeruli analyzed further in (e). **(d)** Heat map showing activity across 227 glomeruli during 150 sec of investigation of the 4 odorants (of 20 min total). Arrowheads indicate responses to carvone and eugenol, shown in expanded views on the right. **(e)** Sensory-driven activity for the two pairs of glomeruli indicated in (c) during investigation of eugenol. Top, proximity to the odor source. Middle, responses of both high- and low-sensitivity pairs. * indicates left-right differences in activity levels across hemispheres; ** indicates recruitment of low-sensitivity pathways close to the odor source. Bottom left, path of the mouse during investigation. Circle sizes indicate strength of sensory responses; X indicates odor source. Bottom right, spatial maps showing mean activity at each location during the entire session, revealing the scale over which odor information is available to the animal.

Next, we used the mesoscope to visualize cellular-resolution activity, a key requirement for characterizing how information is distributed across large neuronal populations. We allowed mice to explore a rectangular arena while imaging the activity of pyramidal neurons through a 3-mm cannula window over area CA1 of hippocampus (Fig. 3a), which plays a central role in spatial navigation as well as learning and memory^6, 22^. Robust single-cell activity was readily apparent throughout the imaging field (Fig. 3b,c; Supplemental Video 2). We extracted intensity time series for each neuron using established analysis packages^23^, yielding signals with high signal-to-noise comparable to established miniscope systems (Fig. 3d,e). The signals from several hundreds of cells could be isolated from even a restricted area of the imaging field (Fig. 3e), and neurons showed spatially modulated firing consistent with other Ca^2+^ imaging studies of CA1^5, 6^.

**Figure 3.**
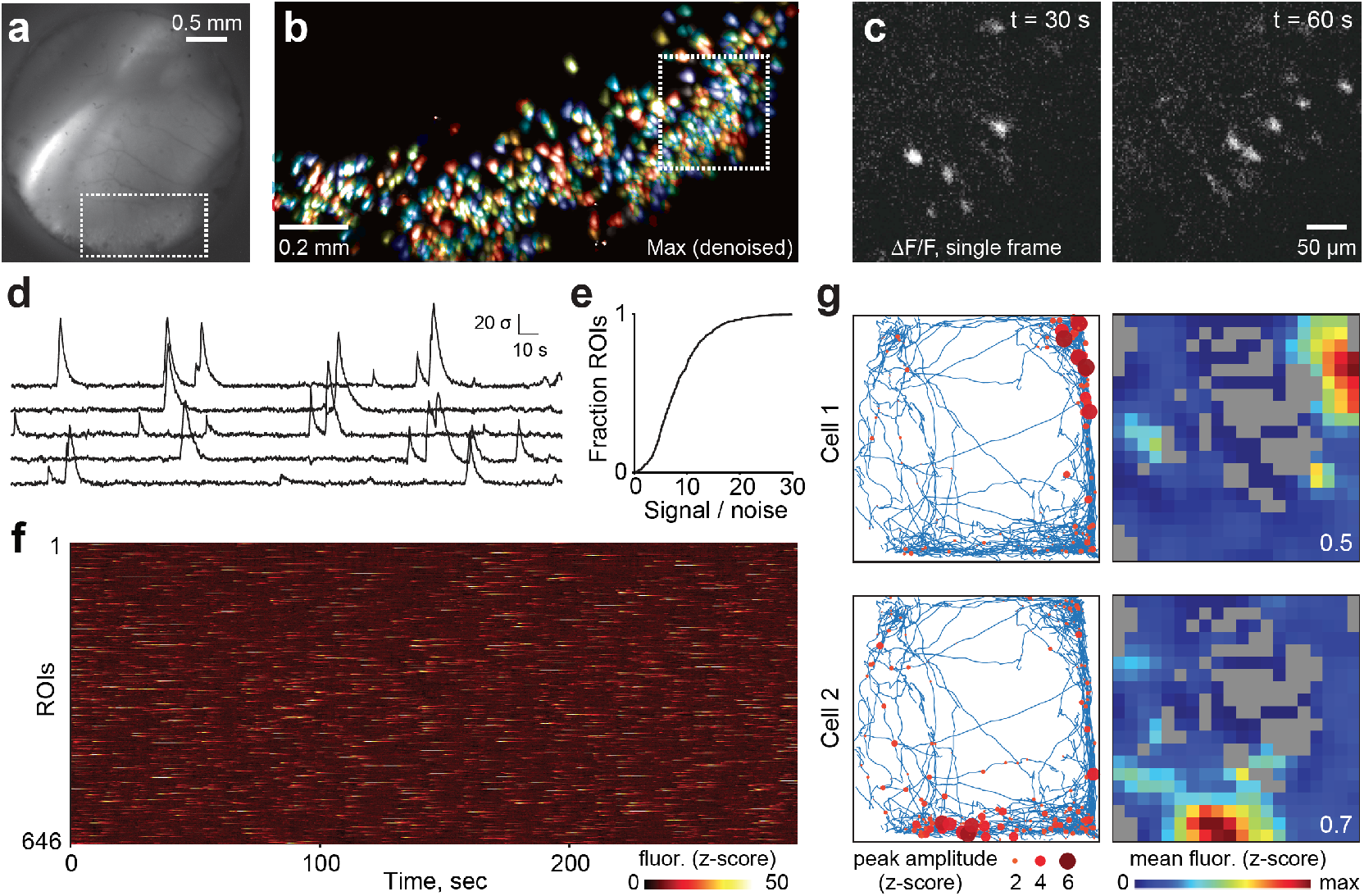
Large-area cellular-resolution imaging of population activity in hippocampal area CA1. **(a)** Raw image frame showing GCaMP expression through the hippocampal cannula window. **(b)** Expanded view of the boxed area in (a), showing a maximum projection of the denoised data; colors show separate ROIs. **(c)** Further enlarged view illustrating single-neuron activity in individual ΔF/F image frames. **(d)** Representative activity traces for five examplar neurons. **(e)** Cumulative histogram of the signal-to-noise ratio for 646 identified ROIs. **(f)** Heat map showing the activity of all neurons imaged in freely moving animals exploring a square arena. **(g)** Locomotor trajectory (left) and heat map (right) showing spatially modulated activity of two example CA1 neurons. Circles reflect location and amplitude of individual Ca^2+^ transients. Colors reflect mean activity averaged within each spatial bin over the session; gray indicates locations not visited by the animal. Numbers indicate the maximum values of the color scale.

Overall, the head-mounted mesoscope provides a powerful new tool for visualizing the widely distributed activity patterns that support diverse behaviors in freely moving animals. While sharing design principles with a system recently described for use in rats^16^, the mesoscope adds several key capabilities. First, it offers the single-cell resolution that is critical for detailed characterization of neural circuit computations. Second, its minimal size and weight allows for chronic implants in mice to take advantage of the extensive genetic tools available in this species. The expanded spatial scales offered by mesoscopic imaging should enable new experiments addressing principles of neural coding and functional interactions between cells within or across brain areas.

## Methods

### Hardware and assembly of the widefield system

Our design builds on the optical design of the open-source ‘FinchScope’ project^3, 4, 9^ as well as the sensor PCB and DAQ hardware developed by the UCLA miniscope group^2, 6^. Optics are assembled in a 3-D printed housing (Formlabs Form2). Blue excitation light from an LED (Luxeon Rebel 470, DigiKey) is collimated by a half ball lens (3mm diameter) and planoconvex lens (6 mm diameter / 10 mm focal length; Edmund Optics # 47-269 and 47-462 respectively) and passed through an excitation filter (470 ± 40 nm, Chroma ET 47/40x, custom diced to 6.25 × 6.25 mm) and dichroic mirror (505 nm shortpass, Thorlabs DMSP505R, 6 × 8.5 mm) onto the brain surface. Emitted fluorescence is reflected horizontally along the optical path by the dichroic, where it passes through a series of 6 mm diameter lenses arranged in a double-Gaussian configuration giving a total magnification of 1.3X (designed in Zemax OpticStudio; planoconvex FL = 18 mm; biconcave FL = 12 mm; Edmund Optics # 47-464 and 48-693). A second short-pass dichroic reflects imaging signals through an emission filter (525 ± 50 nm, Chroma ET 525/50m, 6.25 × 6.25 mm) onto the CMOS sensor. Cellular-resolution imaging was performed with a 5 megapixel Ximea MU9PM-MBRD (2592 × 1944 pixels at 2.2 μm pitch; frame rates of ~5-10 Hz), which offers direct USB sensor output to the acquisition PC. Widefield imaging of MOB dynamics was performed using a custom sensor PCB and data acquisition board developed by the UCLA miniscope group^24^ (ON Semiconductor Python 500, 742 × 480 pixels at 4.8 μm pitch, Miniscope CMOS PCB, LabMaker). The high-power LED excitation needed to illuminate our large imaging areas produced substantial heating, We attached a miniature aluminum heatsink (Adafruit # 1515) to the top of the LED to help dissipate the heating produced by the high-power excitation. We characterized temperature variation at several points along the mesoscope housing, which showed substantial heating at the LED site but little effect at the site of the implant (mean temperature measured at base of implant, Supplementary Figure 3). The total weight of the fully assembled system is approximately 4.2 - 4.5 g.

### Mice, neural labeling, and surgical procedures

All animal experiments were conducted in compliance with the National Institutes of Health Guide for Care and Use of Laboratory Animals, and were approved by the Boston University Institutional Animal Care and Use Committee (protocol # 201800540). Mice were obtained from Charles River (C57Bl6) and the Jackson Laboratory (Tbet-Cre, Ai148, and Thy1-GCaMP6f; stock #’s 024507, 030328, and 024276 respectively) and maintained on a 12/12 hr light cycle in standard cages with *ad lib* access to food and water. Mice were group housed prior to implants, and singly housed afterwards to prevent damage to the implants.

Sensory maps in main olfactory bulb (see Fig. 2) were imaged in Tbet-Cre::Ai148 mice where GCaMP6f is selectively expressed in second-order neurons in main olfactory bulb^25^, avoiding the non-uniformity typical of viral approaches on the 3-4 mm scales of the mesoscope field of view. Adult mice of both sexes were anesthetized with isoflurane (1.5%, diluted in pure O2 delivered at 1.2 L/min) and placed in a stereotax for surgical procedures. We used intact-skull windows constructed by thinning the bone over the dorsal bulbs to a thickness of ~100 μm and covering with cyanoacrylate (Vetbond, TK), optical adhesive (Norland type 81), and a 4mm glass coverslip (Warner Instruments).

Hippocampal imaging (see Fig. 3) was performed in 8-12 week old animals, either in C57BL/6 mice obtained from Charles River or in Thy1-GCaMP6f transgenics line 5.17 obtained from Jackson Labs (ref [^26^]; data not shown). For viral labeling, wildtype animals were anesthetized with isoflurane, a burr hole was made in the skull above the target area, and the CA1 region was stereotaxically injected with 250 nL of virus (AAV9-Syn-GCaMP6f.WPRE.SV40, University of Pennsylvania Vector Core, titer ~6e12 GC/ml) using a 10 nL syringe (World Precision Instruments, Sarasota, FL) fitted with a 33 gauge needle (NF33BL; World Precision Instruments, Sarasota, FL). Injection coordinates were AP: −2 mm, ML: 1.4 mm, DV: −1.6 mm. Virus was injected at a rate of 40 μl/min controlled via a microsyringe pump (UltraMicroPump3–4; World Precision Instruments, Sarasota, FL). Upon recovery, mice underwent surgery for a custom imaging window where the overlying cortical tissue above the hippocampus was carefully aspirated and the cannula window was placed above the CA1 injection site. The cannula window consisted of a stainless steel tube (OD 3.17 mm, ID 2.36 mm, height 2 mm) adhered to a circular coverslip (size 0; OD 3 mm) using a UV-curable optical adhesive (Norland Products). The bottom of the cannula window was attached with Kwik-Sil (WPI) and the tube was attached to the skull with dental acrylic (Metabond, Parkell Inc.). For cortical imaging, a cranial window ~3mm in diameter was opened over posterior cortex, filled with sterile 1% low melting point agarose, and covered with a 4 mm diameter coverslip which was fastened with cyanoacrylate and dental acrylic (Metabond, Parkell). Mice were allowed to recover for 4-6 days after procedures, and supplied with trimethoprim via drinking water to help maintain clarity of hippocampal and cortical windows. After recovery, mice were briefly re-anesthetized with isoflurane and placed in the stereotax for attachment of the mesoscope, which was positioned while imaging using an XYZ micromanipulator to determine the correct focal plane, and attached over the window implants using dental acrylic.

### Resolution testing

Zemax OpticStudio was used to model the expected optical transfer through the system at the center of the imaging plane (the optical axis) and one millimeter lateral to the center (Fig. 1a). The transfer function is expressed as the normalized amplitude modulation of gratings with varying spatial frequencies (line pairs/mm, fig.1d) in two perpendicular orientations. These expected transfer functions were confirmed by placing grating targets (‘airforce’ target in fig 1d left, 100% modulation between dark and bright lines) at either the center point or 1mm laterally (red and blue in fig 1d). The transfer modulation was measured in the image captured by the frame grabber in both orientations (bold and dashed lines in fig. 1d) as *M* = (*I_max_ - I_min_) / (I_max_ + I_min_*), where *I_max_* and *I_min_* are the pixel values of the light and dark stripes.

### Ca^2+^ imaging and data analysis

Mice were placed in a square 45 × 45 cm behavioral arena and allowed to move freely during imaging sessions. For olfactory bulb imaging, four sterile swab tips were loaded with odorant compounds (benzaldehyde, (+)-carvone, eugenol, and amyl acetate; 5% dilution in mineral oil, 5 μL each; Sigma-Aldrich), and placed in the corners of the arena. For hippocampal imaging, the four walls of the arena were covered with visually distinct patterns to provide clear navigational cues. The brain was illuminated at 470 nm with total power between ~6-12 mW over a circular area approximately 4.3 mm in diameter. Imaging data were acquired with custom Matlab code that captures simultaneous neural and behavioral video streams with millisecond-precision timestamps. MOB data were acquired at 30 or 60 Hz at lower resolution. Sensory responses were analyzed with custom Matlab code, where raw fluorescence timeseries were converted to ΔF/F by dividing by a ‘baseline’ image averaged over ≥100 frames. MOB data contained both focal signals arising from glomeruli at the MOB surface, as well as a diffuse component due to scattered light from deeper dendrites and somata, which was removed by subtracting a low-pass spatially filtered version of each frame^27^. ROIs corresponding to individual glomeruli were selected manually in ImageJ based on a maximum projection of the full ΔF/F timeseries. Epochs containing sensory responses used for further analysis were identified from simultaneously recorded behavioral video. Data from CA1 were acquired at approximately 7 Hz using an ROI encompassing the entire ~2.5 mm-diameter cannula window with sensor gain set to 12. Non-rigid motion correction was applied to the image series using NoRMCorre^28^, and ROIs were identified and quantified using Caiman^29^. Signal-to-noise ratios for each trace were computed by CaImAn as the negative logit transform of the average probability that calcium events could be attributed to baseline fluctuations. To characterize spatially modulated activity, fluorescence traces were deconvolved and divided by noise values estimated from high frequency spectral power to produce a ‘z-scored’ estimate of neural activity transients. This measure does not directly report neural spiking activity, but allows closer alignment of neural activity onsets with spatial exploration behavior. These Ca^2+^ events were indicated on the behavioral trajectory of the animal, extracted using DeepLabCut^30^, and heat maps showing ‘place’ activity were constructed by averaging these values in 3 cm spatial bins.

## Acknowledgments

We are grateful to David Boas and the BU Neurophotonics Center for valuable advice, design resources, and institutional support. This work was supported by NIH/NINDS award R24 NS098536 to TJG, TMO, and IGD, NARSAD Young Investigator Grant #27202 to AC-M, and an Edmund Optics Educational Award to DPL.

## Author information

### Contributions

DPL contributed to design and implementation of the mesoscope and performed olfactory imaging experiments. IAC contributed design and optimization of the optical imaging path. KAB and JT performed imaging experiments and data analysis. LKW performed imaging experiments. HJG prepared animals for imaging. WWY contributed to design and implementation of the system. YC contributed to characterizing resolution performance. LNP, WAL, KK, ACM, TJG, TMO, and IGD contributed to design and implementation of the mesoscope. KAB and IGD performed data analysis. IGD wrote the manuscript.

### Competing interests

The authors declare no competing interests. TMO is an employee of Facebook Reality Labs.

### Data availability

Data illustrating the performance of the imaging system and findings of this study are available from the corresponding author upon reasonable request.

## Supplementary Information

**Supplementary Table 1.**
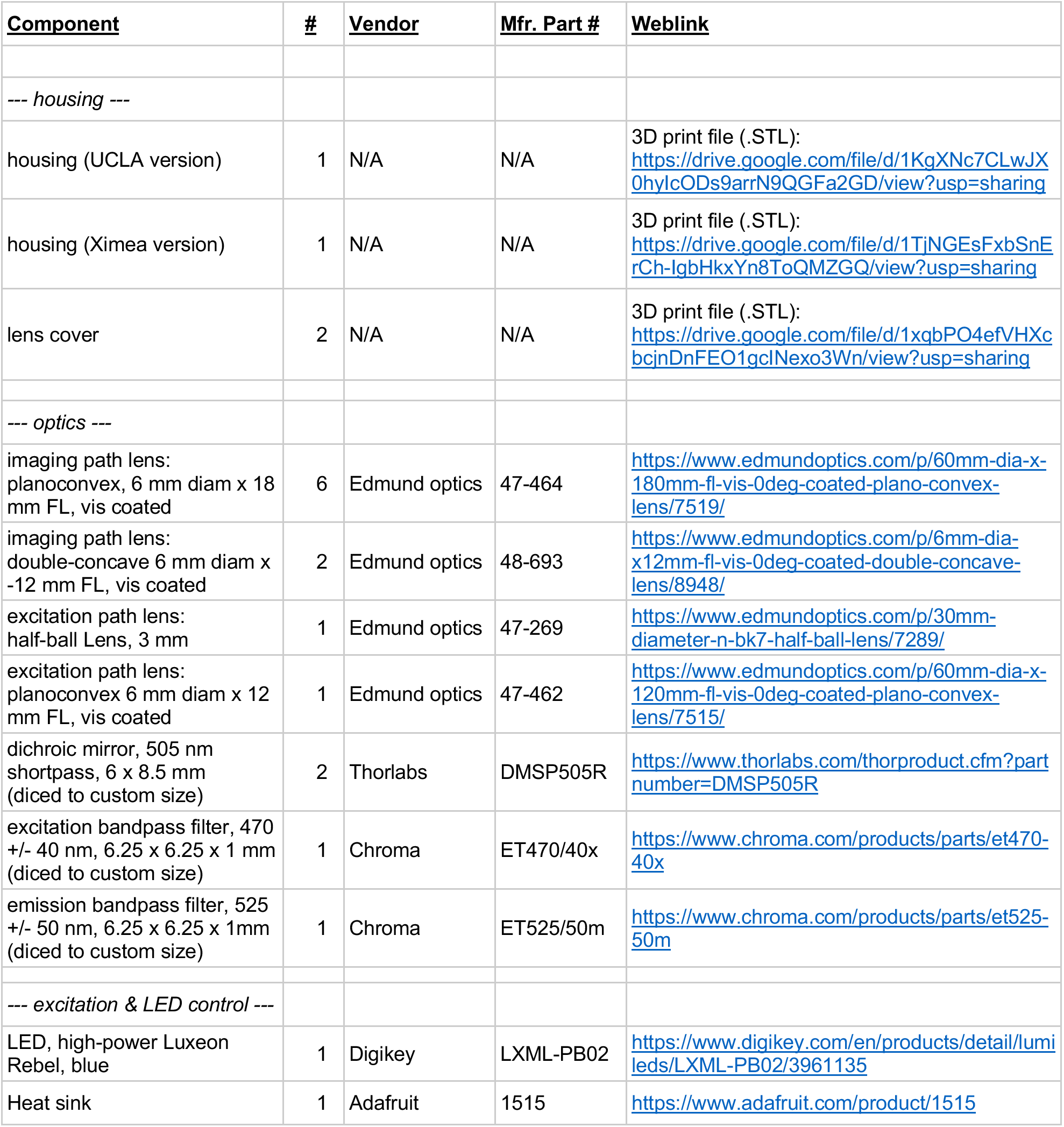
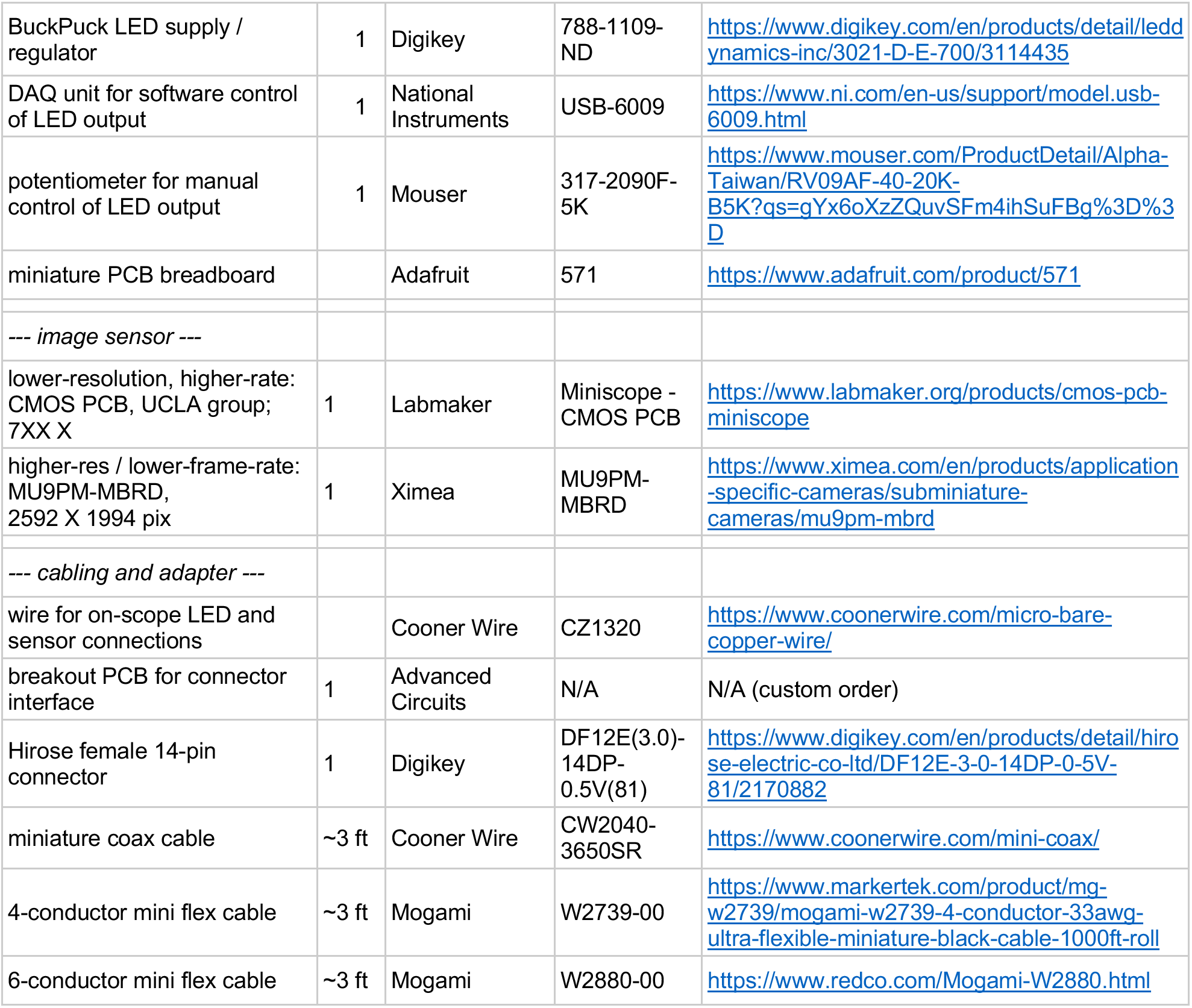
List of components for mesoscope assembly. All components, hardware, and design files needed to build a complete, fully functional system are listed here. An online version of this table is available at: https://docs.google.com/spreadsheets/d/1WQo8Kw7a6SNPagkvxHazFygoOmAfuBclfvQMprrJIXs/edit?ts=5f9b446b#gid=0

**Supplementary Figure 1.**
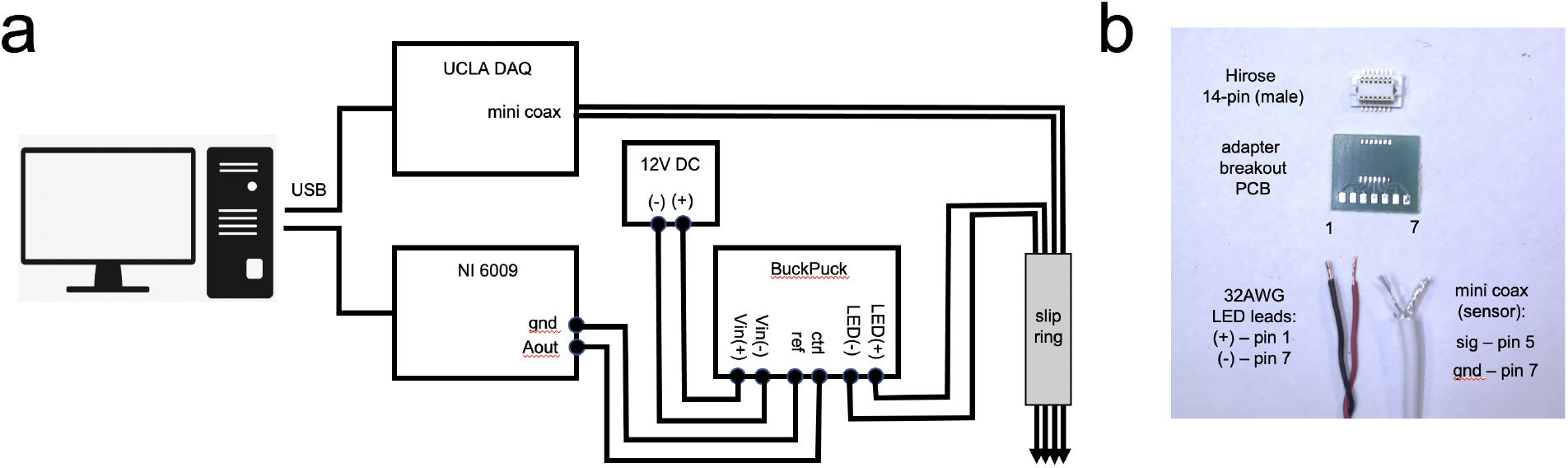
Schematics for connectors and LED control. **(a)** Intensity of the excitation LED is software-controlled using an analog voltage signal generated by a National Instruments USB DAQ board, applied to the control pins of a BuckPuck LED. The circuit also incorporates manual control with a potentiometer and a 3-position switch for selecting between software control, manual control, and off modes. Both sensor and LED signals are routed through a passive 4-channel slip ring to allow unrestricted rotational movement. **(b)** Cables and connectors used to relay image data signals and power for the sensor and excitation LED. Hirose connectors (female, DF12E(3.0)-14DP-0.5V(81); male, DF12A(3.0)-14DS-0.5V(81)) were selected for their balance of small size, light weight, and easy yet stable attachment in implanted animals. Connectors were interfaced with a miniature breakout board (custom design; Advanced Circuits).

**Supplementary Figure 2.**
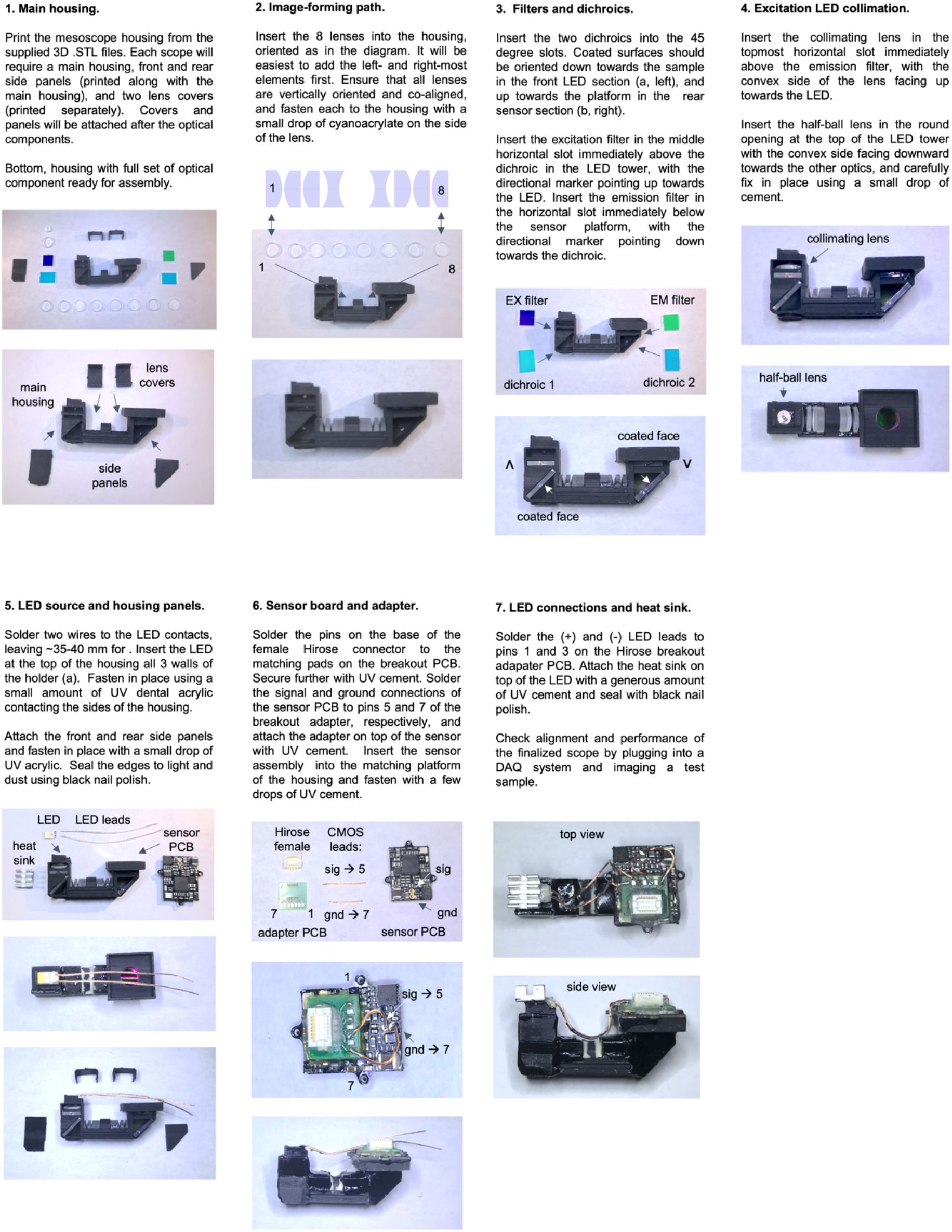
Assembly of the Widescope system.

**Supplementary Figure 3.**
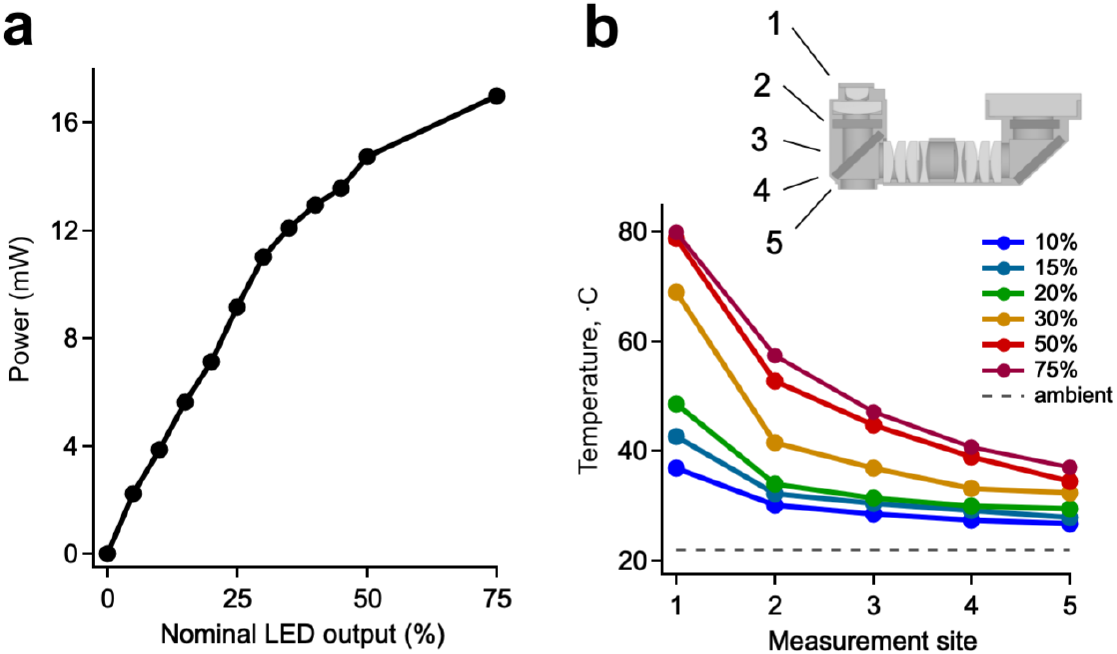
Excitation light intensity and thermal management. **(a)** Total excitation power, delivered to a total area of 16 mm^2^, measured at the imaging plane of the mesoscope as a function of nominal software settings. Imaging sessions typically used a range of 15-20%. LED intensity was set by applying a scaled control voltage spanning the range of the BuckPuck regulator, and excitation power was measured using a commercial power meter (Thorlabs PM100A). Plot shows averaged data from 3 different mesoscopes. **(b)** Temperature measured at the indicated sites on the isolated body of a fully assembled mesoscope using a fine-gauge thermocouple at the sites indicated by the numeric labels. Colors show nominal excitation power levels set by software controls; typical operating ranges are approximately 15-30%. The cement used for implantation adds further mass and surface area for heat dissipation, and temperature at the implant base adjacent to the skull indicated minimal heating (measured at 29.5 and 33.3 °C in two mice).

**Supplementary Figure 4.**
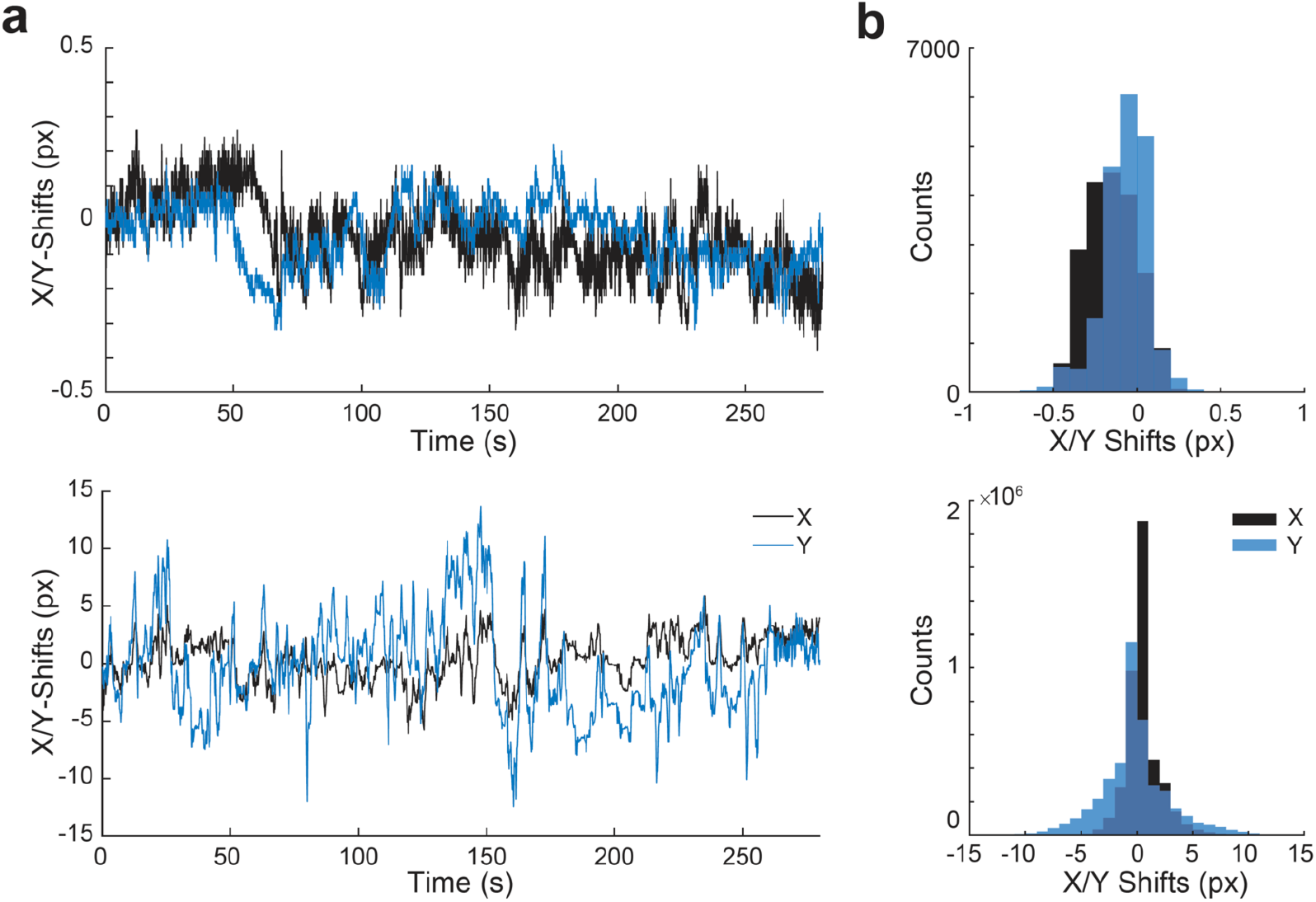
Motion artifacts and image registration. Quantification of image movement during representative recording sessions in freely moving animals. **(a)** Plot of the pixel shifts calculated for registration of a typical image series using NoRMCorre^28^. Top, data from a rigid registration algorithm for olfactory bulb imaging through a thin-skull window. Each pixel corresponds to ~3.8 μm in the image plane. Bottom, parallel data from hippocampal CA1 imaging sessions where each pixel corresponds to ~1.7 μm in the image plane. Values reflect shifts calculated for one of several patches used for a nonrigid algorithm. **(b)** Histogram showing the distribution of shifts applied over longer imaging sessions. Top, olfactory bulb; bottom, CA1.

**Supplementary Video 1.** Dual-hemisphere widefield imaging of sensory responses in the main olfactory bulb during natural investigatory behaviors. Left, investigative behavior of a freely moving mouse within the behavioral arena. Right, corresponding sensory-driven activity in MOB shown as color-coded ΔF/F. Focal signals from surface glomeruli were emphasized by applying a high pass spatial filter (see Methods). Available at: https://drive.google.com/file/d/1yuYPI3LH2DZmFZJcKJKh40Ybqrw9yRdD/view?usp=sharing

**Supplementary Video 2.** Cellular-resolution imaging in hippocampal area CA1. Left panel, exploratory behavior of the mouse within the behavioral arena. Middle, raw fluorescence data collected from the ~3 × 3 mm cannula window. Right, colormapped representation of the ‘denoised’ activity traces assigned to the spatial neuron footprints identified using Caiman. Available at: https://drive.google.com/file/d/1oM3zlqYPoSUvEz0CzDYUAa4GZFoc2CaW/view?usp=sharing

## Notes

https://drive.google.com/file/d/1yuYPI3LH2DZmFZJcKJKh40Ybqrw9yRdD/view?usp=sharing

https://drive.google.com/file/d/1oM3zlqYPoSUvEz0CzDYUAa4GZFoc2CaW/view?usp=sharing

### Competing Interest Statement

The authors have declared no competing interest.

